# Source-sink dynamics could sustain HIV epidemics in rural communities in sub-Saharan Africa: the case of Malawi

**DOI:** 10.1101/468298

**Authors:** Justin T. Okano, Katie Sharp, Laurence Palk, Sally Blower

**Author notes:** These authors contributed equally (Justin T. Okano, Katie Sharp).

## Abstract

**Background:** Approximately 25.5 million individuals are infected with HIV in sub-Saharan Africa (SSA). Epidemics in this region are generalized, show substantial geographic variation in prevalence, and are driven by heterosexual transmission; populations are highly mobile. We propose that generalized HIV epidemics should be viewed as a series of micro-epidemics occurring in multiple connected communities. Using a mathematical model, we test the hypothesis that travel can sustain HIV micro-epidemics in communities where transmission is too low to be self-sustaining. We use Malawi as a case study.

**Methods:** We first conduct a mapping exercise to visualize geographic variation in HIV prevalence and population-level mobility. We construct maps by spatially interpolating georeferenced HIV-testing and mobility data from a nationally representative population-level survey: the 2015-16 Malawi Demographic and Health Survey. To test our hypothesis, we construct a novel HIV epidemic model that includes three transmission pathways: resident-to-resident, visitor-caused and travel-related. The model consists of communities functioning as “sources” and “sinks”. A community is a source if transmission is high enough to be self-sustaining, and a sink if it is not.

**Results:** HIV prevalence ranges from zero to 27%. Mobility is high, 27% of the population took a trip lasting at least a month in the previous year. Prevalence is higher in urban centers than rural areas, but long-duration travel is higher in rural areas than urban centers. We show that a source-community can sustain a micro-epidemic in a sink-community, but only if specific epidemiological and demographic threshold conditions are met. The threshold depends upon the level of transmission in the source- and sink-communities, as well as the relative sizes of the two communities. The larger the source than the sink, the lower transmission in the source-community needs to be for sustainability.

**Discussion:** Our results support our hypothesis, and suggest that it may be rather easy for large urban communities to sustain HIV micro-epidemics in small rural communities; this may be occurring in northern Malawi. Visitor-generated and travel-related transmission may also be sustaining micro-epidemics in rural communities in other SSA countries with highly-mobile populations. It is essential to consider mobility when developing HIV elimination strategies.

## Introduction

The HIV pandemic is concentrated in sub-Saharan Africa (SSA), where ~25.5 million individuals are infected with the virus [1]. Epidemics in SSA countries, unlike those in resource-rich countries, are generalized epidemics driven by heterosexual transmission. The conventional wisdom is that transmission has to be self-sustaining in order to sustain an epidemic. Specifically, one infected individual needs to infect - over their lifetime - at least one other individual. This condition is expressed in the field of transmission modeling in terms of the Basic Reproduction Number, 𝓡_0_; specifically, that 𝓡_0_has to be greater than one [2]. Here, we propose that this is not a necessary condition for HIV epidemics in countries in SSA that have very mobile populations and substantial geographic variability in prevalence. We hypothesize that travel, if it results in viral introductions, could sustain HIV epidemics in communities where transmission is too low to be self-sustaining. We test this source-sink hypothesis by using the epidemic in Malawi as a case study. We first conduct a mapping exercise to determine if there are high levels of, and geographic variability in, HIV prevalence and population-level mobility. We construct these maps using georeferenced HIV-testing and mobility data from a nationally representative population-level survey: the 2015-16 Malawi Demographic and Health Survey (MDHS) [3]. To test our hypothesis, we developed a new conceptual framework for modeling generalized HIV epidemics in SSA. Our framework enables us to model viral introductions that are driven by travel/mobility and includes geographic variation in HIV prevalence and demography. We use this model to analyze the epidemiological interactions between a community that acts as a source and one that acts as a sink; transmission is self-sustaining in the source-community, but not in the sink-community.

Population mobility is very high in SSA due to the necessity of traveling for employment [4-7], this type of travel is often referred to as temporary (or circular) migration. Both men and women spend time away from their families in order to work in agriculture or the textile industry, or as domestic or mine workers. The greatest travel tends to be in rural areas, where poverty is high and there are very few employment opportunities. Results from recent empirical studies indicate that high levels of mobility could be a major driver of HIV transmission in generalized epidemics [4, 5, 8-10]. Migrants and travelers in many SSA countries have been found to engage in riskier behaviors when they are away from home and are more likely to be infected with HIV than non-migrants/travelers. Additionally, several phylogenetic studies of HIV in SSA have shown that in certain communities, strains are being “imported” from other geographic locations [11, 12]; the introduction of viral strains may be due to visitors infecting residents, or residents becoming infected when they travel.

Approximately 85% of the population of Malawi live in rural areas, there are only four cities in the entire country (Figure 1A). The average prevalence of HIV is ~9%, although there is substantial geographic variability in HIV prevalence throughout the country [13, 14]. This variation has been shown to be associated with geographic variation in risk behavior: the number of lifetime sex partners is greater in urban areas (where prevalence is highest) than in rural areas [13]. The recent Migration and Health in Malawi study, which collected longitudinal data from 4,265 men and women living in rural areas, has provided important new insights into the complex relationship between the HIV epidemic and population mobility [15]. Results show that HIV-infected individuals are significantly more likely to be rural-urban, rural-town, or rural-rural migrants than uninfected individuals. Additionally, that HIV-infected migrants are more likely to return to their village of origin than uninfected migrants.

**Figure 1:**
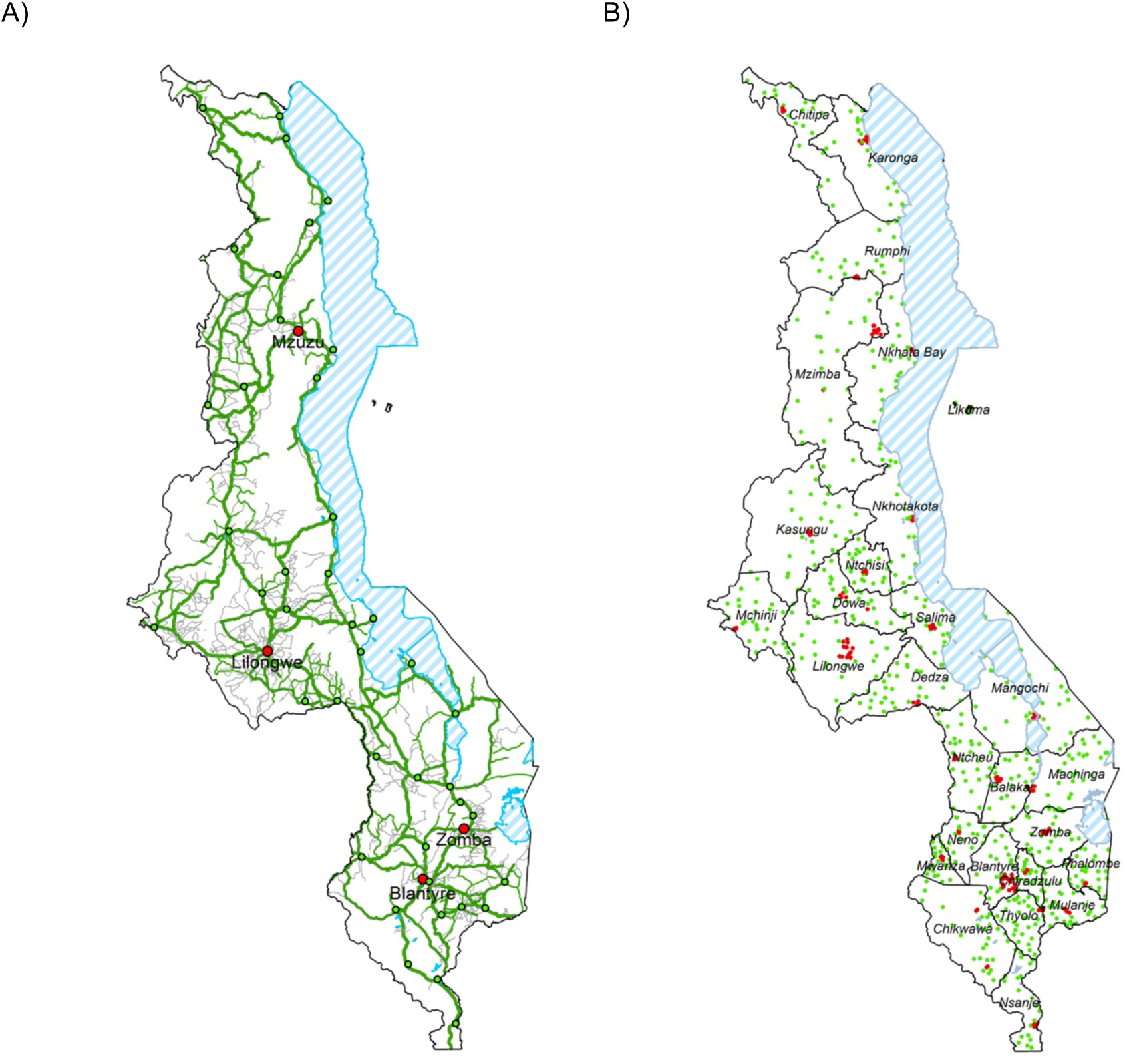
Maps of Malawi. **(A)** Map of Malawi’s road network and cities. Main roads are shown in green, residential roads in gray. Towns are denoted with green circles, the four major cities are denoted with red circles. Lakes are shown by striped blue regions. **(B)** Map showing administrative boundaries for the 28 healthcare districts in Malawi and the sample cluster locations for the 2015-16 MDHS. Locations of clusters in rural locations are shown by green circles, locationof clusters in urban centers by red circles.

Our novel modeling framework considers the transmission dynamics of HIV in a geographically structured population (i.e., a meta-population) where communities have a specific spatial location and function as sources and sinks. We define a source to be a community where transmission amongst residents is self-sustaining [16]; for the micro-epidemic in this community, 𝓡_0_ > 1. We define a sink to be a community where transmission is not self-sustaining [16]; in this community, 𝓡_0_ < 1. To model mobility/travel, we use a generalized Lagrangian approach proposed by Castillo-Chavez and colleagues [17-19]. This modeling approach allows individuals to remain identifiable as residents of their home community when they are in other communities. Notably, our modeling framework is generalizable and can be used to model an HIV epidemic in any country in SSA.

## Methods and Model Analysis

### MDHS data

The 2015-16 MDHS used a two-stage cluster design: 173 urban and 677 rural clusters [3], Figure 1B. The study was designed to collect a representative sample of the population. Demographic, behavioral and mobility data were collected from ~33,000 individuals in ~27,500 households. Additionally, participants who were 15 and older were tested for HIV infection. Each individuals’ test results were linked to their demographic, behavioral and mobility data. The overall response rate was very high: 97% for women, 94% for men. Participation in HIV-testing was also very high: 96% for women, 92% for men. HIV-testing and mobility data were georeferenced based on the location of the sampling cluster.

### Geospatial mapping

We constructed a country-level epidemic surface prevalence (ESP) map to visualize HIV prevalence (as a percentage) in 15-49 year olds [20]. To construct the map, we used data from the 14,010 individuals (aged 15-49 years old) who were tested for HIV in the MDHS, and calculated the proportion who were infected at each cluster site. We then spatially interpolated the cluster-level prevalence data [20]. We used an adaptive bandwidth kernel density estimation method [21]; the kernel density function was modeled as a two-dimensional Gaussian distribution. For smoothing, we chose a ring size of 300 individuals. The R programming package prevR was used to implement this methodology [22].

To construct the mobility maps, we used the individual-level data from the MDHS to calculate the proportion who reported travelling in the previous 12 months at each cluster site. We then employed the same methodology, that we used to construct the ESP map, to spatially interpolate the cluster-level mobility data [20]. The mobility maps show the percentage of 15-49 year olds, at any specific geographic location, that have travelled in the previous 12 months [5, 20]. We constructed two mobility maps: one shows the proportion taking at least one overnight trip in the previous 12 months, the other shows the proportion taking a trip that lasted for a month or more in the previous 12 months.

### An HIV epidemic meta-population model of sources and sinks

The model is designed to represent a generalized epidemic in a country in SSA where the population is highly mobile. The model consists of *n* communities, where *n* specifies the total number of communities in the country. Communities can differ with respect to their demographics (the number of residents), the severity of their micro-epidemic (HIV prevalence), and their connectivity (the proportion of the community that travel to other communities). Mobility/travel is modeled (and hence communities are connected) by specifying a time allocation matrix. Matrix coefficients specify the proportion of time residents spend in their home communities, and the proportion of time they spend in each of the other communities in the meta-population. Due to the movement of individuals, the size of each community can vary over time; at any time, a community is composed of its’ own residents and those from the other communities.

In the model, individuals can become infected with HIV through sexual transmission in their home community or in another community. We model three transmission pathways: (i) resident transmission (individuals acquire HIV in their home community from other residents of their home community), (ii) visitor-caused transmission (individuals acquire HIV in their home community from a resident of another community), and (iii) travel-related transmission (individuals acquire HIV in another community). These three transmission pathways are shown for two communities, that are connected by travel, in Figure 2.

**Figure 2:**
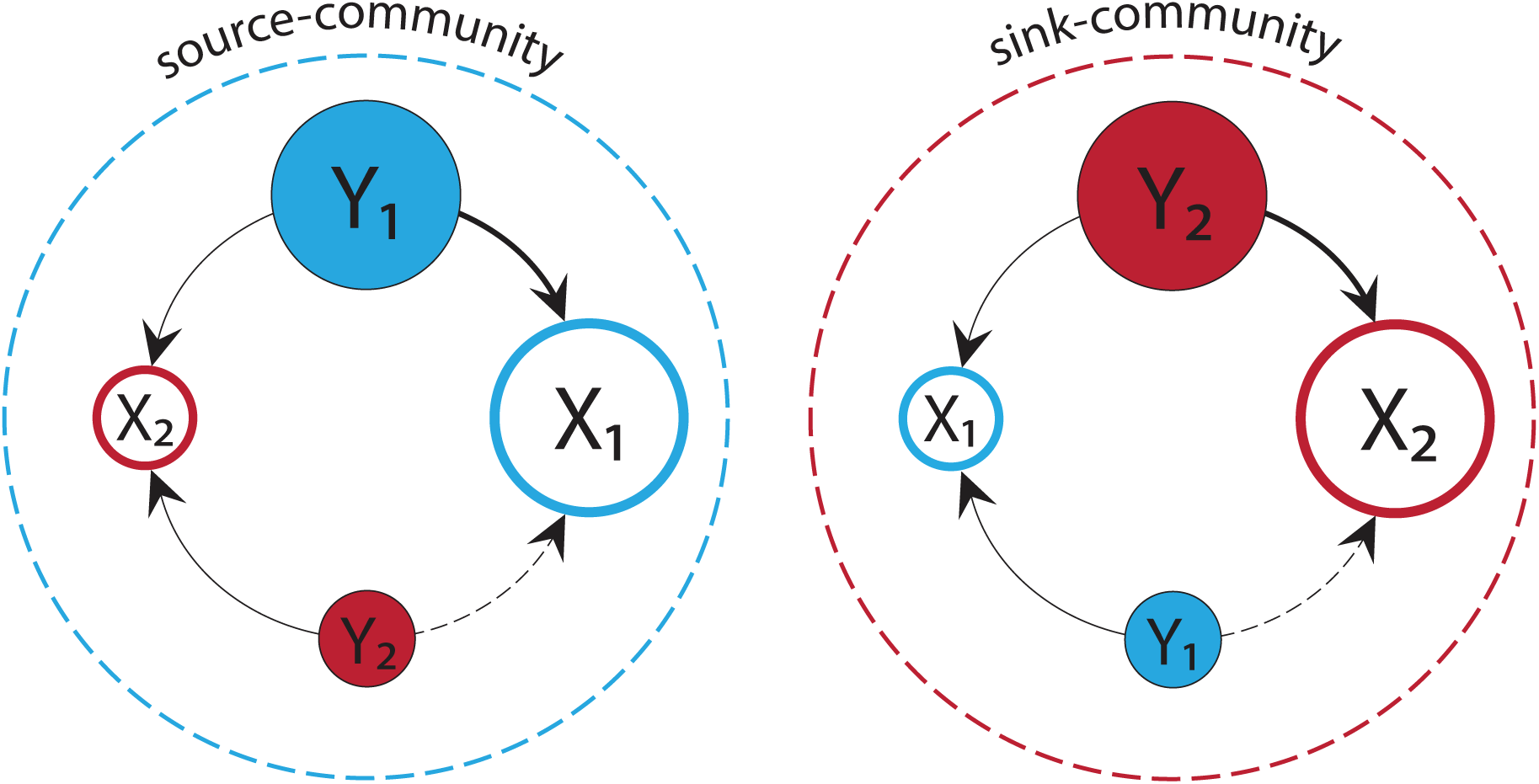
Flow-diagram of two-connected communities in the meta-population model. The diagram shows the three transmission pathways for a source-community (blue dotted circle) and a sink-community (red dotted circle). Blue circles represent residents from the source-community; red circles represent residents from the sink-community. Susceptible residents from each community are denoted by *X_1_* and *X_2_*, respectively. HIV-infected residents from each community are denoted by *Y_1_* and *Y_2_*, respectively. The three transmission pathways are resident-to-resident transmission (arrows with bold lines), visitor-caused transmission (arrows with dotted lines), and travel-related transmission (arrows with thin lines). The two communities are connected due to mobility/travel.

The risk of acquiring HIV is community-specific rather than individual-specific. This could be due to differences in cultural norms and/or opportunities that affect the level of risky behavior amongst the communities. For example, in urban centers individuals have a greater opportunity to have more sex partners than in rural areas. The risk in any community varies with time; this is due to the movement of individuals, changes in the infection status of those who move, changes in community size, changes in prevalence, and increases/decreases in incidence in all of the connected communities in the meta-population.

The meta-population model is specified by the following system of nonlinear differential equations, which determine the rate of change in the number of uninfected (*X_i_*) and infected (*Y_i_*) individuals in community *i:*

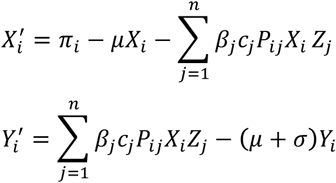

where *π_i_* is the rate at which individuals enter the sexually active class in community *i*, *μ* is the per capita background death rate, *β_j_* is the average probability of transmission per partnership in community *j*, *c_j_* is the average rate of acquisition of sex partners in community *j*, and σ is the per capita death rate due to HIV. *P_ij_* represents the proportion of time that residents of community *i* spend in community *j* and are coefficients in the time allocation matrix. *Z_j_* is the proportion of all individuals (residents and visitors) in community *j* who are infected with HIV and is calculated as
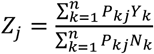 where *N_k_* is the number of residents of community *k*; *N_k_* = *X_k_* + *Y_k_*.

### Defining sources and sinks

To define sources and sinks we analyzed the meta-population model as a system of isolated communities; i.e., we assumed no mobility/travel. In this case, in each community, transmission can only occur amongst residents. The equations for each of the *i* isolated communities are:

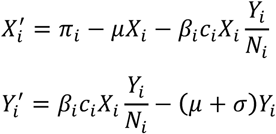

Using these two equations and the next generation matrix methodology [23], we derived the **Isolated-community Reproduction Number**, *Q_i_*. *Q_i_* is defined as the average number of secondary HIV infections that one infected individual generates (over their lifetime) in an isolated community (i.e., in a community where individuals cannot enter or leave), given that all individuals are susceptible.

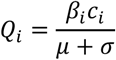

If *Q_i_* > 1, the isolated community is defined as a source. Transmission amongst residents is self-sustaining: it is high enough to maintain a micro-epidemic within the community. If *Q_i_* < 1, the isolated community is defined as a sink. Transmission amongst residents is too low to be self-sustaining: it is too low to maintain a micro-epidemic within the community.

### Modeling two communities that are connected by travel

The equations representing HIV transmission dynamics in two connected communities in the meta-population model are specified by the following four equations:

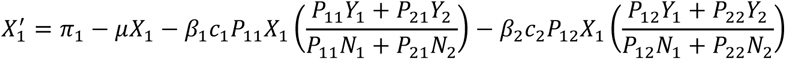

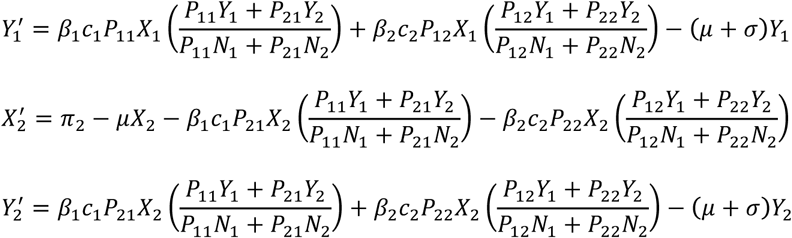

In this case, the time allocation matrix has four coefficients: *P*_12_, *P*_21_, *P*_11_ and *P*_22_. *P*_12_ represents the proportion of time residents of community 1 spend in community 2 on average, *P*_21_ represents the proportion of time residents of community 2 spend in community 1 on average. *P*_11_ and *P*_22_ represent, for each community, the proportion of time residents spend in their home community on average.

Using these four equations and the next generation matrix methodology [23], we derived an analytical expression for 𝓡_0_. 𝓡_0_ is defined to be the average number of secondary HIV infections that one infected individual generates (over their lifetime) in a system of two connected communities, given that all individuals are susceptible.

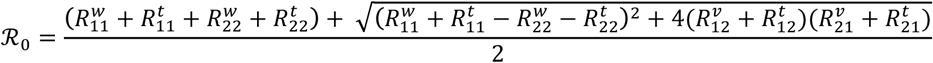

An 𝓡_0_ of greater than one signifies that micro-epidemics are sustained in both communities; the greater the value of 𝓡_0_ above one, the higher the prevalence of HIV. An 𝓡_0_ of less than one signifies that a micro-epidemic only occurs in the source-community (in the case of a connected source- and sink-community), or in neither community (in the case of two connected sink communities).

As can be seen from the analytical expression for 𝓡_0_, it is the root of a quadratic equation. The average number of secondary infections that one infected individual generates (over their lifetime) is the result of a complex interaction of the three transmission pathways in the two connected communities. Each pathway has its own reproduction number: the Connected-community reproduction number, the Visitor-caused reproduction number, and the Travel-related reproduction number. These reproduction numbers are sub-components of 𝓡_0_.

The **Connected-community Reproduction Number** is defined as
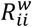 for community *i*. The number represents the average number of secondary HIV infections that are generated by one infected resident (over their lifetime) in the other residents of community *i*, when in their home community. It is the product of the proportion of time residents of community *i* spend in their home community on average, the fraction of susceptible individuals in community *i* who are residents of community *i*, and the isolated-community reproduction number for community *i*.

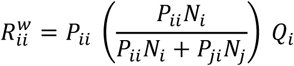

The **Visitor-caused Reproduction Number** is defined as
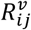 for community *i*. The number represents the average number of secondary HIV infections that are generated by one infected visitor (over their lifetime) in the residents of the community they are visiting. It is the product of the proportion of time residents of community *i* spend in community *j on average*, the fraction of susceptible individuals in community *i* who are residents of community *i* and the isolated-community reproduction number for community *i*.

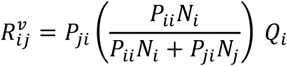

The **Travel-related Reproduction Number** is is defined as
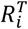 for community *i*. The number represents the average number of secondary infections that occur in individuals of community *i* when in community *j*. The two components of
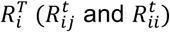 reflect the fact that when an individual spends time in another community they can be infected by residents of that community, as well as by travelers from their home community.

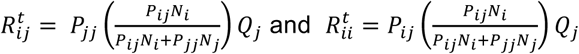

### Testing the hypothesis

Our hypothesis, that viral introductions could sustain HIV epidemics in communities where transmission is too low to be self-sustaining, can be stated mathematically as:

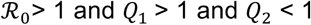

where *Q*_1_ represents the isolated-community reproduction number for the source-community, and *Q*_2_ represents the isolated-community reproduction number for the sink-community.

We tested this hypothesis by conducting an uncertainty analysis, and then performing a threshold analysis. In the uncertainty analysis we calculated values for 𝓡_0_ [24]. We also calculated, for both communities, values for their connected-community reproduction number, their visitor-caused reproduction number, and their travel-related reproduction number. We made these calculations by specifying parameter ranges (for *Q*_1_, *Q*_2_, *P*_12_, and *P*_21_) and using Latin Hypercube sampling [25] to sample each of these ranges 10,000 times. This procedure resulted in 10,000 calculated values for each of the reproduction numbers.

To motivate parameter ranges, we assumed that the source-community would be an urban community, and the sink-community a rural community. For the source-community, we varied *Q*_1_ between 1.10 and 1.50; this translates to a prevalence range for the micro-epidemic in the source-community (when isolated) of 9% to 33%. To translate *Q*_1_ into prevalence, we used the relationship: prevalence = 1 – 1/*Q*_1_ [2]. The range for *Q*_1_ was based on previous analysis of prevalence data from Malawi [13]. We explored the effect of *Q*_1_, the isolated-community reproduction number for the sink-community, over a wide range: 0.50 and 0.95. To set ranges for *P*_12_, and *P*_21_ we made two assumptions. First, we assumed that residents would spend most of their time in their home community. Second, we assumed that residents of the urban community would spend less time in the rural community than residents of the rural community would spend in the urban community. On the basis of these two assumptions we varied the range for the percentage of time that residents from the source-community spend in the sink-community (*P*_12_) between 5% and 15%, and the range for the percentage of time that residents from the sink-community spend in the source-community (*P*_21_) between 10% and 35%. We set the ratio of community size (*N_2_* : *N_1_*) to be 0.80, so that the source was slightly larger than the sink.

We conducted the threshold analysis by using the values from the uncertainty analysis for 𝓡_0_. Specifically, we identified the threshold values of *Q*_1_ and *Q*_2_ at which 𝓡_0_ = 1. At these threshold values, the source-community is just able to sustain a micro-epidemic in the sink-community. We also explored the effect of the relative ratio of community size (sink to source) on the threshold values of *Q*_1_ and *Q*_2_.

## Results

The ESP map shows HIV prevalence (in 15-49 year olds) varies substantially geographically, from 0% to 27% (Figure 3A). Large-scale spatial patterns are apparent: there is a strong north-south trend in increasing prevalence and a substantial urban-rural difference. Prevalence is highest in the four cities in Malawi (Blantyre and Zomba in the Southern region, Lilongwe in the Central region, and Mzuzu in the Northern region) and in the fishing villages along Lake Malawi. Prevalence is lowest in rural areas.

**Figure 3:**
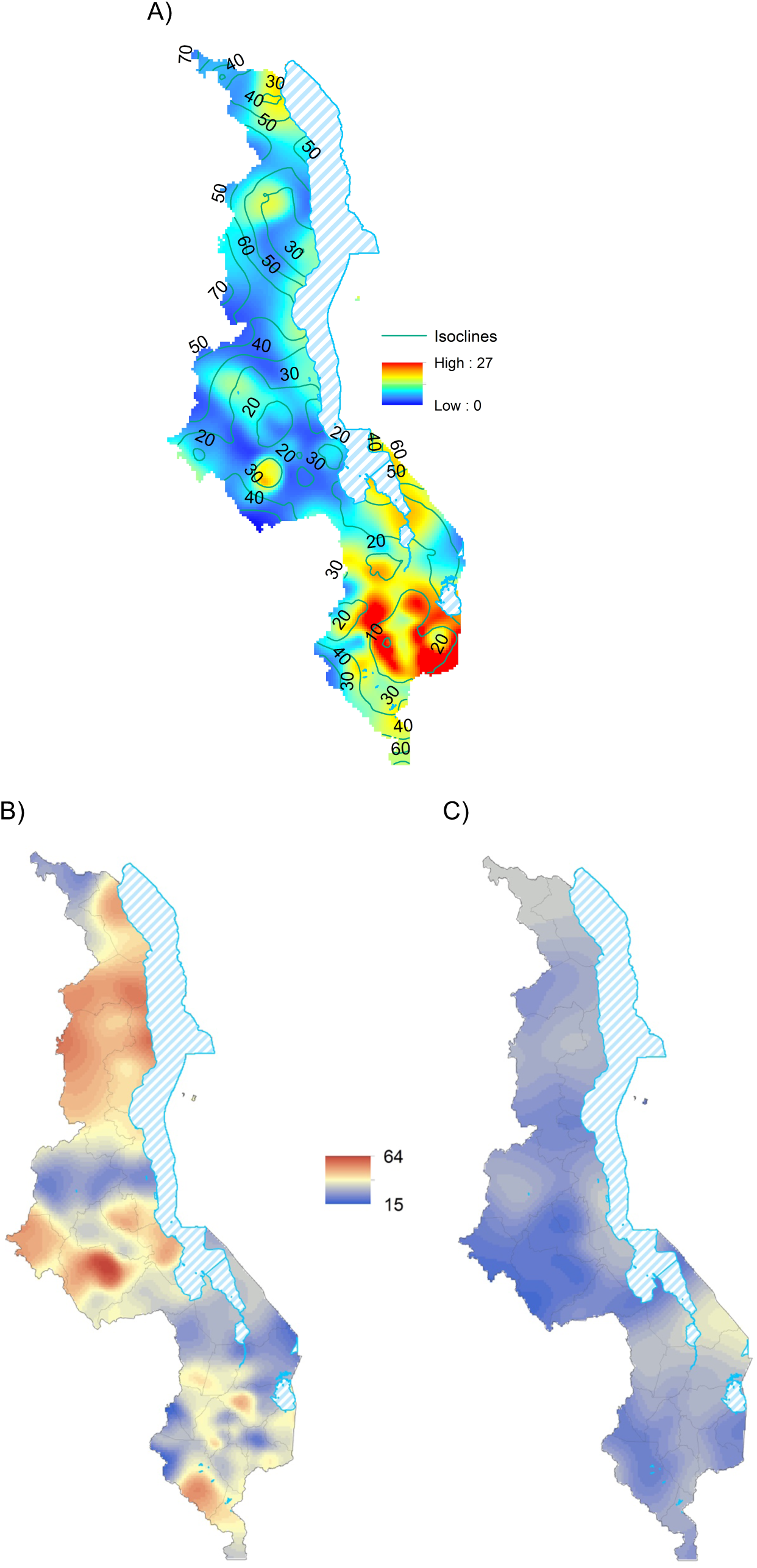
HIV prevalence and mobility maps. **(A)** ESP map for Malawi. Color scale indicates the HIV prevalence (%) in individuals aged 15-49 years old. The isoclines show the size (in km) of the radii of the smoothing circles used in the spatial interpolation to calculate the ESP map; these circles contain 300 individuals between 15 and 49 years old. **(B)** Mobility map showing the percentage of individuals aged 15-49 years old, at each geographic location, who made one or more overnight trips in the previous 12 months. **(C)** Mobility map showing the percentage of individuals aged 15-49 years old who took a trip lasting at least one month in the previous 12 months.

The population of Malawi is highly mobile: 38% of 15-49 year olds took at least one overnight trip in the previous 12 months (Figure 3B), 27% were away for at least a month (Figure 3C). Large scale-spatial patterns are evident; they tend to correspond to urban-rural differences. Almost a third of residents in the Northern region, where HIV prevalence is low and which is predominantly rural, had been away from home for at least a month in the previous 12 months. Notably, ~60% of the residents of the capital city (Lilongwe) had been on at least one overnight trip in the previous 12 months, with the majority of trips lasting less than a month.

At a lower spatial resolution than shown on the ESP map, at the level of the healthcare district, there is still substantial geographic variation in prevalence and mobility/travel (Table 1). HIV prevalence varies from 3% to 21%, the percentage of the population who took at least one overnight trip in the previous 12 months varies from 26% to 56%, and the percentage of the population who were away for at least a month in the previous 12 months varies from 20% to 34%. There is also a high degree of variability in the size of the population amongst districts: ranging from ~1,185,000 in Lilongwe (which includes Lilongwe City) to only ~5,000 in Likoma (Table 1).

**TABLE 1:** Characteristics of healthcare districts in Malawi. The table shows, for each of Malawi’s 28 healthcare districts, the number of individuals (15-49 years old) living in the districts in 2015 (the numbers are projections from the 2008 census data [35]), HIV prevalence among individuals (15-49 years old), the proportion of individuals in this age group who took at least one trip in the previous 12 months, and the proportion of individuals in this age group who took a trip lasting at least one month in the previous 12 months. Mobility and prevalence data are from the 2015-16 MDHS [3].

| District | Population | Prevalence | ≥ 1 trip | ≥ 1 month |
| --- | --- | --- | --- | --- |
| Balaka | 172,269 | 9.1% | 0.33 | 0.31 |
| Blantyre | 653,720 | 18.2% | 0.36 | 0.25 |
| Chikwawa | 239,521 | 7.7% | 0.41 | 0.27 |
| Chiradzulu | 142,426 | 9.4% | 0.32 | 0.33 |
| Chitipa | 92,526 | 3.3% | 0.26 | 0.32 |
| Dedza | 321,350 | 3.3% | 0.35 | 0.23 |
| Dowa | 343,055 | 3.7% | 0.34 | 0.22 |
| Karonga | 147,835 | 10.2% | 0.47 | 0.34 |
| Kasungu | 362,742 | 4.3% | 0.27 | 0.29 |
| Likoma | 5,048 | 7.6% | 0.38 | 0.22 |
| Lilongwe | 1,184,696 | 7.3% | 0.56 | 0.20 |
| Machinga | 257,994 | 6.4% | 0.29 | 0.33 |
| Mangochi | 439,559 | 10.5% | 0.31 | 0.29 |
| Mchinji | 257,987 | 5.3% | 0.48 | 0.22 |
| Mulanje | 254,656 | 20.9% | 0.32 | 0.26 |
| Mwanza | 46,939 | 7.7% | 0.26 | 0.20 |
| Mzimba | 511,961 | 3.6% | 0.47 | 0.29 |
| Neno | 67,949 | 11.0% | 0.33 | 0.30 |
| Nkhata Bay | 122,937 | 6.5% | 0.49 | 0.29 |
| Nkhotakota | 166,913 | 7.4% | 0.34 | 0.30 |
| Nsanje | 120,656 | 11.5% | 0.36 | 0.29 |
| Ntcheu | 245,511 | 8.2% | 0.30 | 0.34 |
| Ntchisi | 121,657 | 4.7% | 0.45 | 0.22 |
| Phalombe | 161,031 | 16.3% | 0.36 | 0.24 |
| Rumphi | 95,851 | 6.4% | 0.52 | 0.28 |
| Salima | 180,971 | 3.1% | 0.45 | 0.31 |
| Thyolo | 289,421 | 12.2% | 0.36 | 0.28 |
| Zomba | 362,506 | 13.6% | 0.40 | 0.29 |

The 10,000 values of 𝓡_0_ calculated in the uncertainty analysis are shown in the form of a histogram in Figure 4A. The median value of 𝓡_0_ is 1.13, with an interquartile range (IQR) of 1.06-1.21. Notably, 92% percent of the values of 𝓡_0_ are greater than one, signifying that the source-community is capable of sustaining a micro-epidemic in the sink-community. These results provide strong support for our hypothesis that viral introductions could sustain HIV epidemics in communities where transmission is too low to be self-sustaining. The results also show that the source-community is not always able to sustain a micro-epidemic in the sink-community; this is the case when the calculated value for 𝓡_0_ is less than one. Therefore, Figure 4A demonstrates the existence of a sustainability threshold: i.e., that there are specific conditions that need to be met in order for a source-community to be able to sustain a micro-epidemic in a sink-community.

**Figure 4:**
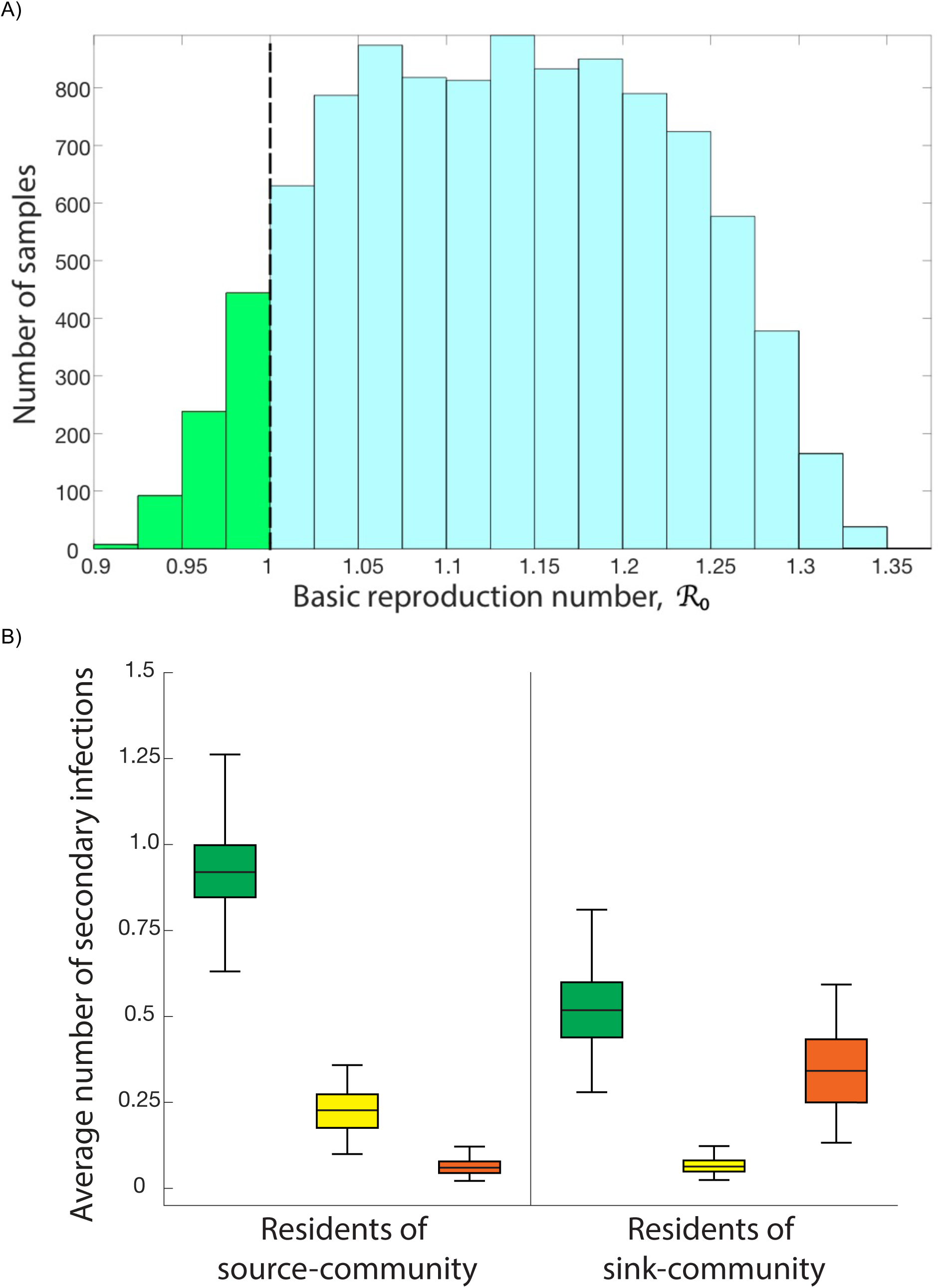
Results from the uncertainty analysis. **(A)** Frequency distribution of 𝓡_0_. Values are calculated from 10,000 samples of parameter ranges using Latin Hypercube Sampling. Green bars show the samples where the source-community is not able to sustain a micro-epidemic in the sink-community, light blue bars show the samples where the source-community can sustain a micro-epidemic in the sink-community. The dotted black line denotes the sustainability threshold at 𝓡_0_ = 1. **(B)** Boxplots, for the samples where 𝓡_0_ > 1, show the average number of secondary infections caused by one infectious individual (over their lifetime) for each of six reproduction numbers:
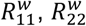 (green data),
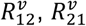 (yellow data),
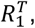 and
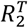 (orange data).

Figure 4B provides a mechanistic understanding of the complexity of the transmission dynamics that occur when a source-community sustains a micro-epidemic in a sink-community. The boxplots show the values calculated in the uncertainty analysis for the two communities for their connected-community reproduction number, their visitor-caused reproduction number, and their travel-related reproduction number. The y-axis shows the average number of secondary infections generated by one individual, over their lifetime, for each reproduction number. In both communities, the largest reproduction number is for transmission between residents (green boxplots). Not surprisingly, on average, the number is much higher for the source-community than the sink-community: 0.89 (median) versus 0.50 (median). Because the risk of acquiring HIV is greatest in the source-community, the average number of infections caused by a visitor to the source-community is greater than the average number caused by a visitor to the sink-community (yellow boxplots: 0.22 (median) versus 0.06 (median)). For the same reason, on average, the travel-related reproduction number is greater for a resident of the sink-community than for a resident of the source-community (orange boxplots: 0.35 (median) versus 0.06 (median)).

Figure 5A shows a response hypersurface [26] that has been constructed based on the results of the uncertainty analysis; the response variable is 𝓡_0_. The y-axis shows the range in the values of the reproduction number for an isolated source-community (*Q*_1_); the x-axis shows the range in the values of the reproduction number for an isolated sink-community (*Q*_2_). The black line shows the sustainability threshold at which 𝓡_0_ equals one and the source-community is just able to sustain an epidemic in the sink community. These results show that our hypothesis that viral introductions can maintain HIV micro-epidemics in communities where transmission is too low to be self-sustaining holds true, but only if certain conditions are met. Sustainability is possible in the parameter space above the threshold, but is not possible in the parameter space below the threshold.

**Figure 5:**
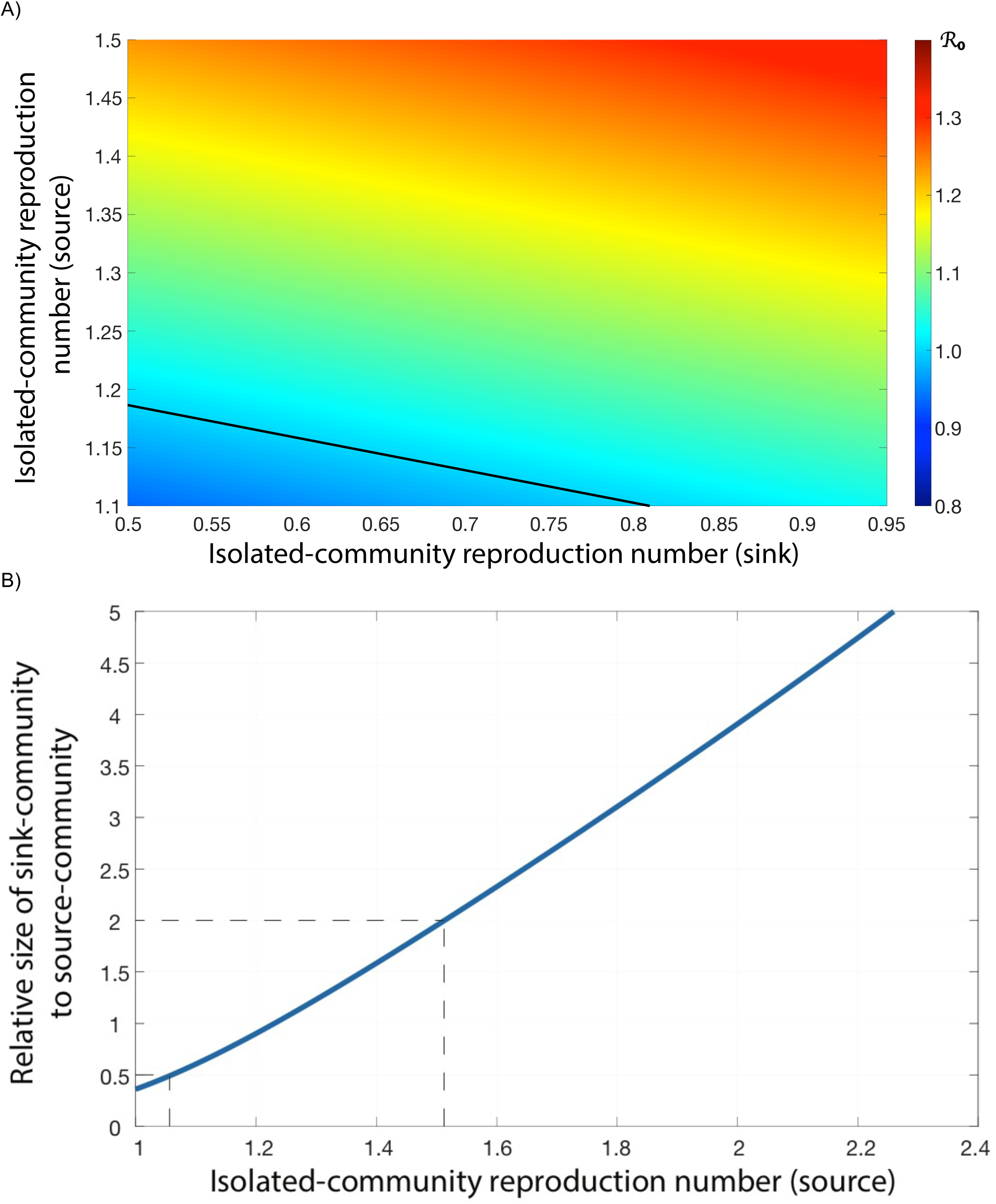
Results from the threshold analysis. Results are based on the 10,000 values of 𝓡_0_ that are calculated in the uncertainty analysis and the ranges for *Q*_1_and *Q*_2_ that are used as inputs in the uncertainty analysis. **(A)** Response hypersurface plot showing the effect of *Q*_1_(y-axis) and effect of *Q*_2_ (x-axis) on the value of 𝓡_0_. The black line shows the sustainability threshold conditions at which 𝓡_0_ = 1. The sink-community is 80% of the size of the source-community. **(B)** The effect of the size of the sink-community relative to the size of the source-community (y-axis) on the sustainability threshold, when *Q*_2_ = 0.50. The blue line shows the threshold values for *Q*_1_ that are necessary to sustain a micro-epidemic in the sink-community.

Figure 5A shows that a source-community can sustain a micro-epidemic in a sink-community even if transmission in the sink-community (which is reflected in the value of *Q*_2_) is extremely low. Sustainability is possible even if one infected individual in the isolated sink-community only generates 0.50 secondary infections over their lifetime. With this low level of transmission, to ensure sustainability, one infected resident of the source-community would need to generate at least 1.19 secondary infections over their lifetime; this means that HIV prevalence in the isolated source-community would need to be at least 16%. Not surprisingly, the higher the transmission in the sink-community, the lower the transmission in the source-community needs to be to exceed the sustainability threshold. If *Q*_2_ equals 0.81, *Q*_1_ only needs to exceed 1.10 (which translates to a prevalence of 9.9% in the isolated source-community) to ensure sustainability, Figure 5A.

Figure 5B shows the effect of the ratio of community size (y-axis) on the minimium (i.e., threshold) value of *Q*_1_ needed to ensure that a source can sustain a micro-epidemic in a sink: *Q*_2_ is held constant at 0.50. The blue line shows the sustainability threshold values for *Q*_1_. The sink is greater than the source when the ratio is greater than one; the source is greater than the sink when the ratio is less than one. The larger the source than the sink, the lower the prevalence in the isolated source-community needs to be for sustainability. Conversely, the smaller the source than the sink, the higher the prevalence in the isolated source-community needs to be for sustainability. Notably if the source is twice the size of the sink, prevalence in the isolated source-community only needs to be greater than 5.7% to ensure sutainability. However, if the sink is twice the size of the source, prevalence needs to exceed 34%.

## Discussion

Our mapping studies show that mobility/travel is high throughout Malawi, and there is a high degree of geographic variation in both prevalence and mobility/travel. The spatial patterns that we have uncovered show that prevalence is higher in urban centers than in rural areas, but that long-duration travel is higher in rural areas than in urban centers. We have presented a new framework, based on considering communities as sources and sinks, for modeling generalized HIV epidemics in SSA. Our model includes geographic variation in HIV prevalence and demography, as well as three transmission pathways: resident-to-resident, visitor-caused, and travel-related. This framework enables us to model viral introductions that are driven by travel/mobility and to test the source-sink hypothesis. We found that - contrary to conventional wisdom - an epidemic can be sustained in a community where transmission is too low to be self-sustaining. We identified the existence of a threshold that shows that sustainability is only possible under certain epidemiological and demographic conditions. We also found that if the size of the source-community is much larger than that of the sink-community, prevalence in the source-community does not have to be very high to ensure sustainability. Our results imply that it would be rather easy for a large urban community to sustain a micro-epidemic in a small rural community.

Our results have significant implications for the control of HIV in Malawi. In order to design effective control strategies, it is essential to understand the underlying transmission dynamics of the generalized epidemic. Our results suggest that Malawi’s epidemic is more complex than currently assumed. In rural areas in the north, communities are small and widely dispersed, the population is highly mobile, travel is of long duration, and HIV prevalence is low. In the one major urban center in the region, the city of Muzu, prevalence is extremely high. It is possible that Muzu is functioning as a source and that transmission is being sustained in the surrounding rural communities as a result of urban-rural and rural-urban travel. If so, reducing within-community transmission without also preventing visitor-caused and travel-related transmission is unlikely to be effective in reducing the number of new infections in northern Malawi. Blantyre, Zomba, Lilongwe and fishing villages along Lake Malawi may also be important sources and be sustaining micro-epidemics in the surrounding rural communities. Phylogenetic analysis could be used, as in Uganda [11, 12] and South Africa [27], to differentiate between localized and imported strains and determine (for specific communities) where transmission is occurring. Such analyses would provide important information regarding where to locate interventions and what type of intervention would be most effective; e.g., pre-exposure prophylaxis or voluntary mass circumcision.

There are several limitations to our study, with respect to both the MDHS data and our modeling framework. The MDHS data are not detailed enough to be able to determine which regions of the country are interconnected through travel, or the spatio-temporal dynamics of mobility patterns. With more detailed datasets, such as those based on mobile phone call records, large-scale network-based analyses of mobility could be conducted. Such analyses, if linked to HIV-testing and demographic data, could provide additional insights into the effect of mobility/travel on HIV epidemics. In our modeling, we have made the reasonable assumption that an urban community is connected by travel to a nearby rural community. We have only used the mobility data to assess the level of population mobility and geographic variation. We have not used these data to parameterize our model. Our modeling framework can be applied to an entire country, but such an exercise would be only worthwhile if the necessary parameters can be estimated and travel patterns that link communities can be determined. We have conducted a parsimonious analysis of a system based on two linked communities in order to gain an initial understanding of a very complex system and to test the source-sink hypothesis. To the best of our knowledge, our model is the first to examine the potential impact of population-level imobility on generalized epidemics in SSA; it is also the first model that includes visitor-caused and travel-related transmission. As such, it is a conceptual advance and provides new insights into an important component of the HIV pandemic. Our current research focuses on a detailed analysis of the meta-population model, using several connected source- and sink-communities. In this more complex system, it is possible for one source-community to sustain micro-epidemics in several sink-communities and, conversely, for multiple source-communities to sustain a micro-epidemic in one sink-community. We are also expanding the modeling framework in order to include gender differences, both in terms of prevalence levels (prevalence is higher in women, than in men, in Malawi [13]) and in time allocation matrices.

Our results have significant implications for the control of HIV epidemics in highly mobile populations throughout SSA. All of the HIV epidemics in this region are generalized and show substantial geographic variation in prevalence. Geospatial models have been used to predict the future of these epidemics, and to develop control strategies based on the allocation of limited resources [28-32]. These models include geographic variation in prevalence, but none of them include mobility; therefore, none of them include visitor-caused or travel-related transmission. Our results highlight the necessity of including mobility when modeling HIV epidemics in SSA. The model, presented in this paper, could be used as a foundation for building more realistic models. This approach would result in more accurate predictions and the design of more cost-effective control strategies for HIV epidemics in SSA. We recommend collecting large datasets that can provide a realistic representation of population-level mobility patterns and their spatio-temporal dynamics [33]; these representations could then be integrated within a modeling framework. High levels of mobility are proving to be a substantial obstacle in the global attempt to eliminate malaria [34]; we suggest that they may also be a substantial obstacle in the global attempt to eliminate HIV.

## Abbreviations

ESP: Epidemic surface prevalence;
IQR: Interquartile range
MDHS: Malawi demographic and health survey;
SSA: sub-Saharan Africa

## Authors’ contributions

JTO, KS, LP, and SB developed the concept and study design, analyzed and interpreted the data and drafted the manuscript. JTO and KS conducted the statistical analyses and model coding. JTO and LP conducted the geospatial analyses. SB supervised the project and wrote the manuscript. All authors read and approved the final manuscript.

## Acknowledgements

The authors are grateful to Carlos Castillo-Chavez for ideas and discussions throughout the course of this research. National Institute of Health Awards R01 AI116493 supported this work.

